# The overlapping modular organization of human brain functional networks across the adult lifespan

**DOI:** 10.1101/2021.11.12.468371

**Authors:** Yue Gu, Liangfang Li, Yining Zhang, Junji Ma, Chenfan Yang, Yu Xiao, Ni Shu, Cam-CAN, Ying Lin, Zhengjia Dai

## Abstract

Previous lifespan studies have demonstrated that the brain functional modular organization would change along with the adult lifespan. Yet, they assumed mutual exclusion among functional modules, ignoring convergent evidence for the existence of modular overlap. To reveal how age affects the overlapping functional modular organization, this study applied a detection algorithm requiring no prior knowledge of the resting-state fMRI data of a healthy cohort (N = 570, 18-88 years). Age-related regression analyses found a linear decrease in the overlapping modularity and the similarity of modular structure and overlapping node (i.e., region involved in multiple modules) distribution. The number of overlapping nodes increased with age, but the increment was distributed unevenly. In addition, across the adult lifespan and within each age group, the nodal overlapping probability consistently exhibited positive correlations with both functional gradient and flexibility. Further, we showed that the influence of age on memory-related cognitive performance might be explained by the change in the overlapping functional modular organization. Together, our results revealed age-related decreased segregation from the perspective of brain functional overlapping modular organization, providing new insight into the adult lifespan change in brain function and its influence on cognitive performance.

## 1. Introduction

Lifespan research has become a hot spot with the intensification of global aging (Bookheimer et al., 2019). In neuroscience, mapping how the human brain network (i.e., human connectome) changes across the lifespan can enhance our understanding of neurocognitive development and decline [for review, see (Zuo et al., 2017)]. Specifically, previous studies on brain functional networks have found the inverted-U trajectories of the local efficiency over the lifespan (Cao et al., 2014), and age-related linearly increase in the shortest-path length and the average clustering coefficient (Sala-Llonch et al., 2014). Additionally, the nodal betweenness was found to decrease in the frontal lobe and occipital lobe while the nodal degree and the nodal efficiency increased in the posterior frontal lobe and parietal lobe over the lifespan (Chandan et al., 2018).

Modules, which are derived from a decomposition of the network, are subcomponents that are internally strongly coupled but externally weakly coupled. As an important topological characteristic of the human brain functional network, the functional modular organization has been widely studied (Alexander-Bloch et al., 2010; Bordier, Nicolini, Forcellini, & Bifone, 2018; Calhoun, Kiehl, & Pearlson, 2008; Jones et al., 2012; Liao, Cao, Xia, & He, 2017; Meunier, Achard, Morcom, & Bullmore, 2009; Meunier, Lambiotte, & Bullmore, 2010). Three essential features of the functional modular organization have been revealed: 1) the identification of modules has shown high reproducibility in each individual (Guo et al., 2012); 2 each module corresponds to specific cognitive performance (Sadaghiani & Kleinschmidt, 2013), such as the visual networks (VIS) supports visual perception (Van Den Heuvel & Pol, 2010) and the fronto-parietal networks (FPN) is involved in initiating and adjusting control (Dosenbach, Fair, Cohen, Schlaggar, & Petersen, 2008); 3) modularity may promote brain adaptation and increase flexibility in response to a changing environment [for review, see (Sporns & Betzel, 2016)] and prevent catastrophic forgetting (Ellefsen, Mouret, & Clune, 2015). Recently, several studies have been focusing on the age-related changes in the brain functional modular organization and suggested that brain segregation reduces as age increases Specifically, lower modularity with weaker intra-module functional connectivity but stronger inter-module functional connectivity was continuously observed across the lifespan (Chan, Park, Savalia, Petersen, & Wig, 2014; Geerligs, Renken, Saliasi, Maurits, & Lorist, 2014; Spreng, Stevens, Viviano, & Schacter, 2016). The age-accompanied functional modular changes were mainly located in the default mode network (DMN), dorsal attention network (DAN), VIS, and FPN (Betzel et al., 2014; Cassady, Ruitenberg, Reuter-Lorenz, Tommerdahl, & Seidler, 2020; Ferreira & Busatto, 2013; Grady, Sarraf, Saverino, & Campbell, 2016; Puxeddu et al., 2020; Spreng et al., 2016). Additionally, these age-related changes of the functional modular structure were found associated with cognitive control and attention performance (Betzel et al., 2014).

Currently, the module detection methods used in the above age-related studies mainly focused on non-overlapping modules, that is, each brain region only belongs to a single module. However, neuroimaging studies have suggested that the human brain functional network is more likely to bear an overlapping modular organization (Lin et al., 2018; Najafi, Mcmenamin, Simon, & Pessoa, 2016; Yeo, Krienen, Chee, & Buckner, 2014), in which a brain region can participate in more than one functional module. Previous studies not only provided rich evidence to support the existence of overlap among functional modules (Bassett et al., 2011; Bullmore & Sporns, 2009; Fries, 2005; Power, Schlaggar, Lessov-Schlaggar, & Petersen, 2013), but also implied that the concept of overlap may pave the way to interpret the flexible and variable relationships between the human brain and cognitive functions in a more realistic manner (Lin et al., 2018; Yeo et al., 2015). In particular, it would be of great interest to study how the overlapping functional modular structure, including the overlapping modules and the overlapping nodes, change across the adult lifespan and how such change induces and/or affects the cognitive functions. However, to date, few studies have examined the changes in overlapping functional modules across the adult lifespan.

To address this issue, we employed resting-state functional magnetic resonance imaging (R-fMRI) to explore the overlapping modular organization of the human brain functional network in 570 healthy participants across the adult lifespan (18-88 years). Firstly, the maximal-clique based multiobjective evolutionary algorithm (MCMOEA; Wen et al., 2016; Lin et al., 2018) was used to identify the overlapping brain functional modular structure of each participant. Secondly, based on the overlapping modules detected, we respectively examined the changing trajectories of the overlapping modules and the overlapping nodes during adult lifespan by regression models and age-based group comparisons. Then, we revealed the functional features of the nodal overlapping probability through functional gradient and flexibility analyses. Finally, we examined how the characteristics of overlapping modules and nodes were related to fluid intelligence and the Benton face recognition test performance, both of which could effectively measure individual memory capacity and were already found sensitive to age (Feng et al., 2020; Kievit et al., 2014).

## 2. Materials and Methods

### 2.1 Participants

Data of 649 participants [age range 18-88 years; mean = 59.24, standard deviation (SD) = 18.55] were obtained from the second stage of the Cambridge Centre for Ageing and Neuroscience (Cam-CAN) (http://www.cam-can.org, Shafto et al., 2014). Among these participants, 79 participants were excluded for having one of the following issues: missing data, image artifacts, and excess head motion. Thus, a final sample of 570 participants (age range 18-88 years, mean = 52.88, SD = 18.44, 287 females) was included in our main analyses. Ethical approval was obtained from the Cambridge shire Research Ethics Committee and all participants gave their written informed consent prior to participation.

### 2.2 Data acquisition and preprocessing

The MRI data were collected on a 3T Siemens TIM Trio System, with a 32-channel head coil. Resting-state fMRI (R-fMRI) data were obtained using an echo-planar imaging sequence parameters: repetition time (TR)/echo time (TE) = 1970 ms/30 ms, flip angle = 78°, number of slices = 32, slice thickness = 3.7 mm, voxel size = 3 mm × 3 mm × 4.44 mm, field of view (FOV) =192 mm × 192 mm and total volumes = 261. The 3D T1-weighted structural images were acquired using Magnetization Prepared Rapid Acquisition Gradient-Echo pulse sequences. The sequence parameters were 1 × 1 × 1 mm^3^ resolution, TR/TE = 2250 ms/2.99 ms, inversion time (TI) = 900 ms, flip angle = 9°, and FOV = 256 × 240 × 192 mm^3^. More details of the data collection can be found in Shafto et al. (2014).

Image preprocessing for R-fMRI was carried out using the Data Processing Assistant for Resting-State fMRI (DPARSF) (http://rfmri.org/DPARSF; Yan, & Zang, 2010) toolbox and SPM8 (http://www.fil.ion.ucl.ac.uk/spm). For each participant, the first six volumes were discarded. The realignment was performed after slice timing to the first volume to correct head motion. Sixty-nine participants were excluded for excess head motion (more than 2 mm or 2°). Then the T1-weighted image was coregistered to the mean functional image after motion correction and then segmented into gray matter, white matter and cerebrospinal fluid tissue images. The head motion corrected functional images were further spatially normalized to the Montreal Neurological Institute (MNI) space using the parameters estimated from T1 unified segmentation (Ashburner & Friston, 2005) and were resampled into 3‐mm isotropic voxels. Finally, the normalized functional images were detrended, regressed out the nuisance variables (Friston’s 24 head motion parameters, global signal, white matter, and cerebrospinal fluid signals) and temporal band-pass filtered to 0.01-0.08 Hz.

### 2.3 Construction of brain functional networks

The brain functional network construction was carried out using the graph theoretical network analysis [GRETNA, http://www.nitrc.org/projects/gretna/, Wang et al., 2015)]. For each participant, we parcellated the whole brain into 264 regions/nodes (Power, Cohen, Nelson, Wig, & Petersen, 2011). Then, we computed the Pearson correlation coefficient between time series of each pair of nodes and generated a 264 by 264 symmetric correlation matrix. Considering the ambiguous biological explanation of negative correlations (Fox, Zhang, Snyder, & Raichle, 2009; Murphy, Birn, Handwerker, Jones, & Bandettini, 2009), we only preserved positive correlations and set the negative correlations as zeros. A binary and undirected functional network was then constructed by thresholding the matrix with 15% sparsity.

### 2.4 Detection of overlapping modules

To detect individual overlapping modules in the brain functional network, the maximal-clique-based multiobjective evolutionary algorithm (MCMOEA) (Lin et al., 2018; Wen et al., 2016) was used. Without assuming the number of overlapping modules, this algorithm evolves a population of candidate overlapping modular structures through customized operators to achieve the optimal tradeoff between two objectives: maximizing the intra-link density whilst minimizing the linter-link density of modules. In this study, the MCMOEA was applied and the application procedure was similar to that introduced by Lin et al. (2018).

### 2.5 Analyses of overlapping modules and overlapping nodes

We derived three measures to capture the characteristics of overlapping modules at the individual level: (1) the number of overlapping modules; (2) the overlapping modularity score; (3) the modular similarity. To capture the characteristics of overlapping nodes, we firstly delineated the spatial pattern of overlapping nodes by visualizing the distribution of the nodal overlapping probability. Then, four individual-level measures were calculated based on the overlapping nodes: (1) the number of overlapping nodes; (2) the membership diversity of overlapping nodes, which was reflected by the numbers of nodes participating in *k* modules (*k* ≥ 2); (3) the modular overlapping percentage regarding the ten classic non-overlapping functional modules specified by Power et al. (2011); (4) the variability in the spatial locations of overlapping nodes with other participants.

Based on the above measures, we then analyzed the effect of age on the overlapping modular structure in two ways. Firstly, using age as the independent variable, each of the seven individual-level measures as the dependent variable and gender as the covariate. Secondly, to perform statistical comparisons to evaluate the age-related changes in overlapping modular organization, we divided the participants into three age groups (Young: 18-45 years, 208 participants; Middle: 46-64 years, 169 participants; Old: 65-88 years, 193 participants) to quantify the between-group differences in the above measures.

To further explore whether and how the functional roles of overlapping nodes were influenced by age, we computed the Pearson correlation coefficients between the nodal overlapping probabilities and two functional indicators (gradient and functional flexibility). Finally, to assess whether and how the brain functional modular organization was related to the cognitive performance (i.e., fluid intelligence and Benton face recognition test) during the adult lifespan, we computed the Pearson correlation coefficients between cognitive performance and each of the above individual-level measures regarding overlapping modules/overlapping nodes. To further explore whether the individual-level overlapping module or node characteristics mediated the age effects on cognitive performance, a mediation analysis was performed.

## 3. Results

### 3.1 Adult lifespan changes of overlapping modules

Among the three measures for characterizing the overlapping modules, a significant influence of age was found on the overlapping modularity (Figure 1A-B) and the modular similarity (Figure 1C-D), but not on the number of overlapping modules. In detail, the overlapping modularity showed negative correlation with age (*R* = −0.204, *P* < 10^−3^; Figure 1A). The overlapping modularity within the elderly group was significantly lower than that of the other two age groups (Old < Young: *P* = 0.002; Old < Middle: *P* < 10^−3^; Figure 1B). These results implied that the functional segregation capability of the human brain gently decreased across the adult lifespan. The modular structure similarity calculated in the context of the entire population was found negatively correlated with age (*R* = −0.417, *P* < 10^−3^; Figure 1C). In addition, the modular structure similarity calculated within each age group coincided with the above decreasing trend, and the permutation test confirmed that the differences of the modular structure similarity between age groups were all significant (Young > Middle > Old: all *P* < 10^−3^; Figure 1D). These results suggested that the overlapping modular structure of the human brain functional network was getting more individual variability as age increased.

**Figure 1.**
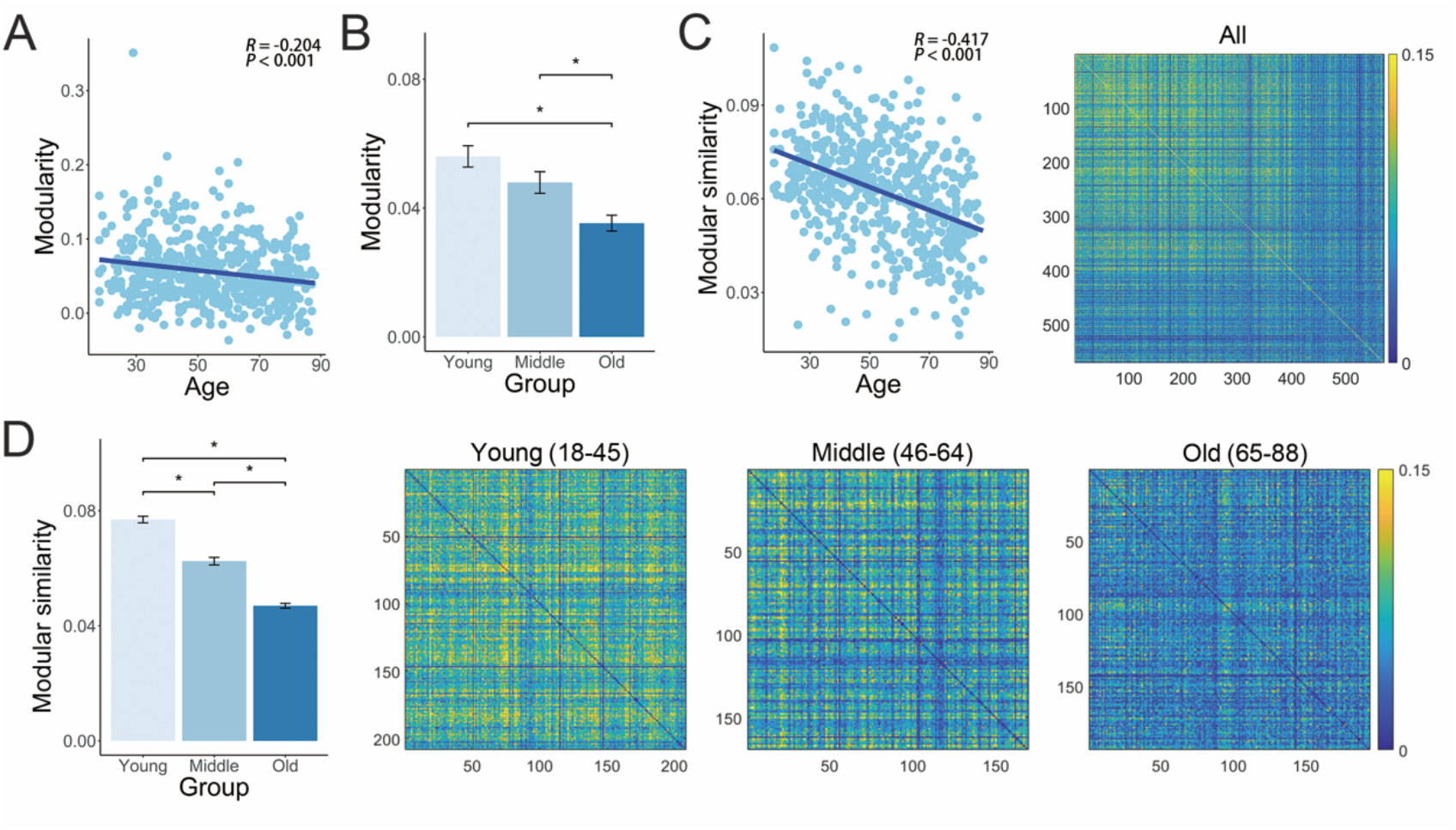
Lifespan changes in overlapping modules regarding (A-B) modularity and its between-group comparison, (C-D) modular similarity and its between-group comparison. In bar plot, the asterisk indicates significant between-group difference *(P* < 0.05, 10000 permutations, Bonferroni-corrected).

### 3.2 Adult lifespan changes of overlapping nodes

Figure 2 depicted the distribution of nodal overlapping probabilities calculated from the entire population of participants and each of the three age groups. Overall, the nodal overlapping probability shown in Figure 2A. The similar distribution patterns were also observed in the three age groups (Figure 2B-D).

**Figure 2.**
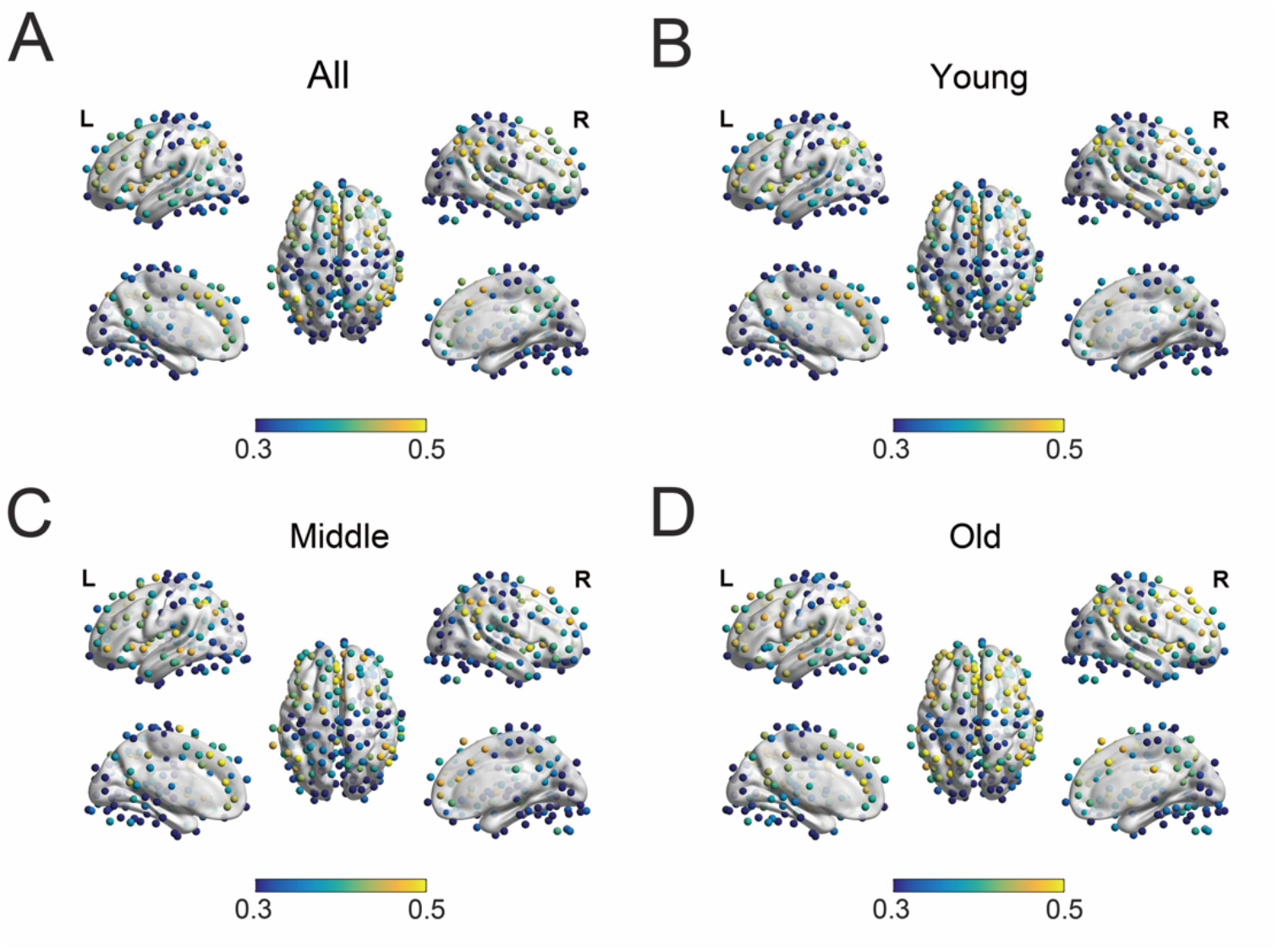
Distribution of nodal overlapping probabilities (A) over all the participants and (B-D) within each of the three age groups.

The number of overlapping nodes linearly increased along adult lifespan (*R* = 0.149, *P* < 10^−3^, Figure 3A), particularly for overlapping nodes that participated in two and three overlapping modules (*k* = 2: *R* = 0.142, *P* = 0.001; *k* = 3: *R* = 0.131, *P* = 0.004; Figure 3B), but not for the overlapping nodes participating in four or more modules (*k* ≥ 4). Between-group comparisons further confirmed that the old group had significantly more overlapping nodes (Old > Young and Old > Middle: all *P* < 10^−3^; Figure 3C) and significantly higher membership diversity (*k* = 2 or 3) (Old > Young and Old > Middle: all *P* ≤ 0.006; Figure 3D) than each of the other two age groups. Additionally, although the overlapping nodes were found distributed in all the ten classic non-overlapping modules, DMN had the largest modular overlapping percentage considered either across the entire adult lifespan or within each age group (Figure 4A-B). We found that the modular overlapping percentage linearly increased with age in VIS (*R* = 0.148, *P* = 0.002), but linearly decreased in FPN (*R* = −0.129, *P* = 0.008; Figure 4C). Between-group comparison analyses also showed that compared with the old group, the modular overlapping percentage of the young group was significantly lower in VIS (*P* < 10^−3^) but remained higher in FPN with a marginally significant difference (*P* = 0.020, Bonferroni-corrected, Figure 4D). Further, Figure 5 showed the regions whose overlapping probabilities exhibited significant between-group differences. Moreover, the individual variability in the spatial pattern of overlapping nodes decreased as age increased (*R* = −0.119, *P* = 0.005; Figure 6A), and significant differences were detected between the old group and each of the other two groups (Old < Young: *P* < 10^−3^, Old < Middle: *P* = 0.002; Figure 6B).

**Figure 3.**
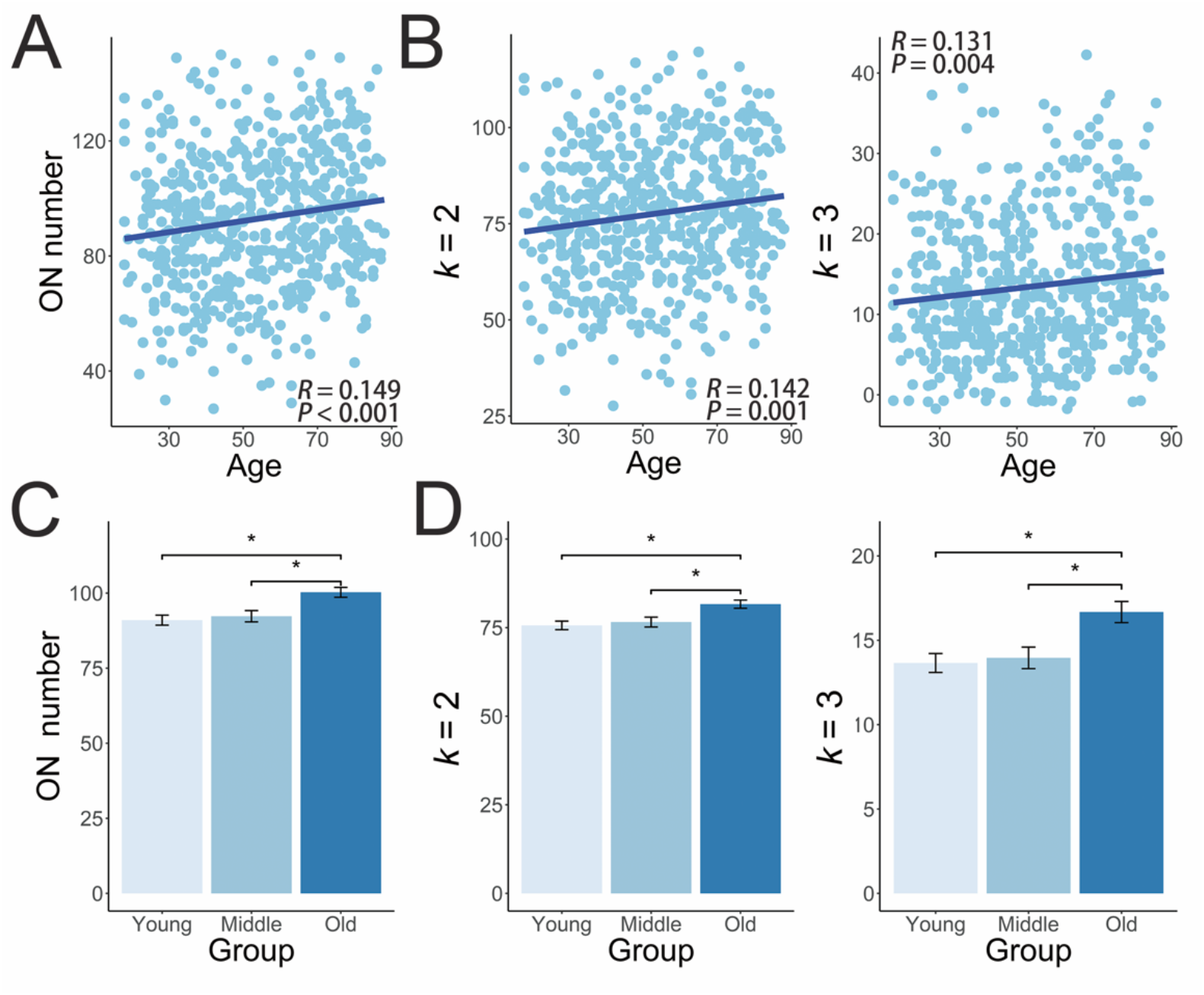
Lifespan changes in overlapping nodes (ON) regarding (A) number, (B) the member ship diversity, the between-group comparison of (C) number and (D) the membership diversity. The light blue nodes denote participants, and dark blue lines denote the aging regression line for linear model. In bar plot, the asterisk indicates significant between-group difference *(P* < 0.05, 10000 permutations, Bonferroni-corrected).

**Figure 4.**
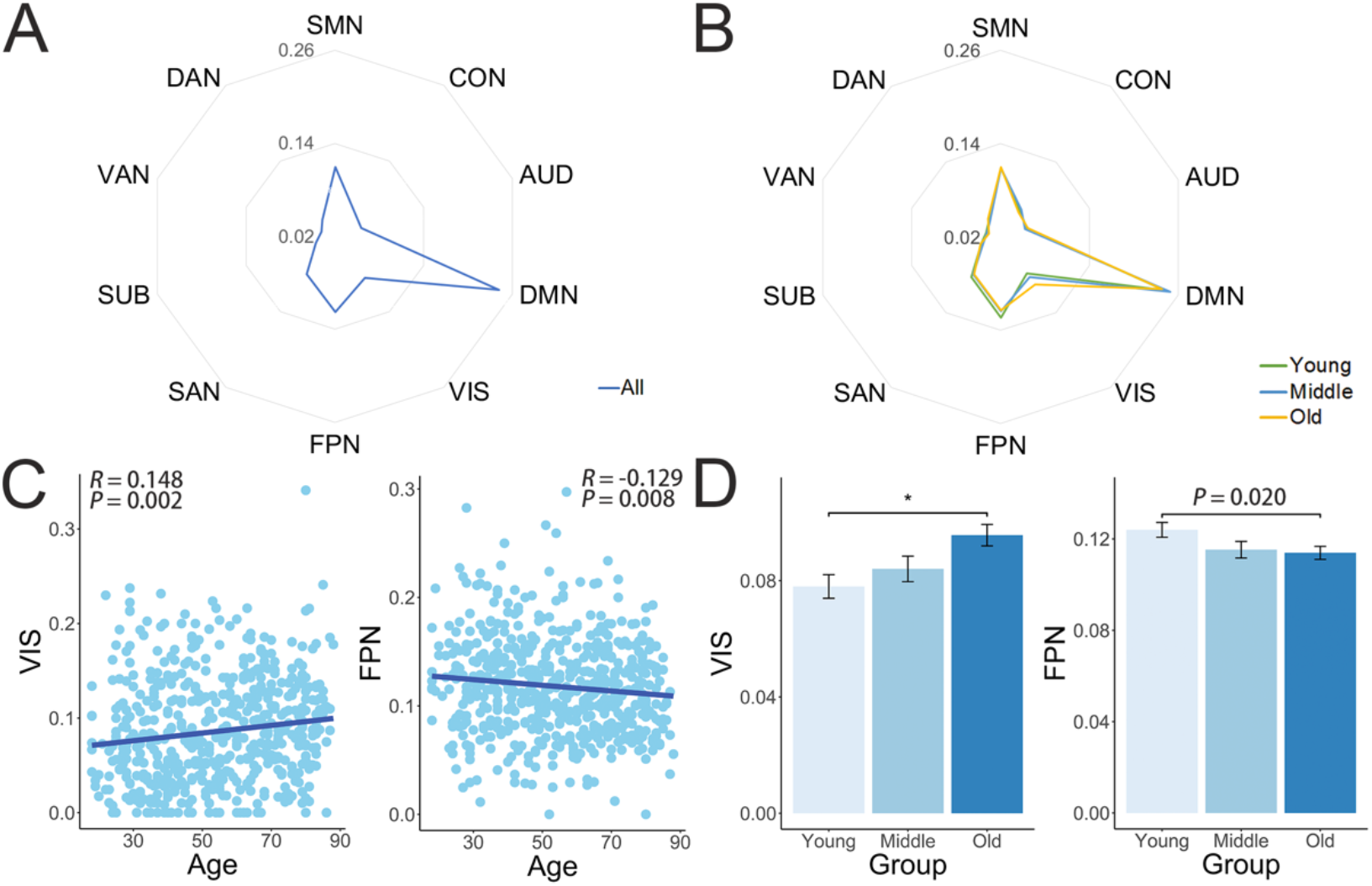
Lifespan changes in overlapping nodes (ON) regarding (A-B) the distribution in ten classic non-overlapp ing modules, (C) the modular overlapping node percentage and (D) its between-group comparison. The light blue nodes denote participants, and dark blue lines denote the aging regression line for linear model. In bar plot, the asterisk indicates significant between-group difference *(P* < 0.05, 10000 permutations, Bonferroni-corrected). Abbreviations: FPN, fronto-parietal network; CON, cingulo-opercular network; SAN, salient network; DAN, dorsal attent ion network; VAN, ventral attention network; DMN, default mode network; SMN, somatosensory-motor network; AUD, audial network; VIS, visual network; SUB, subcortical network.

**Figure 5.**
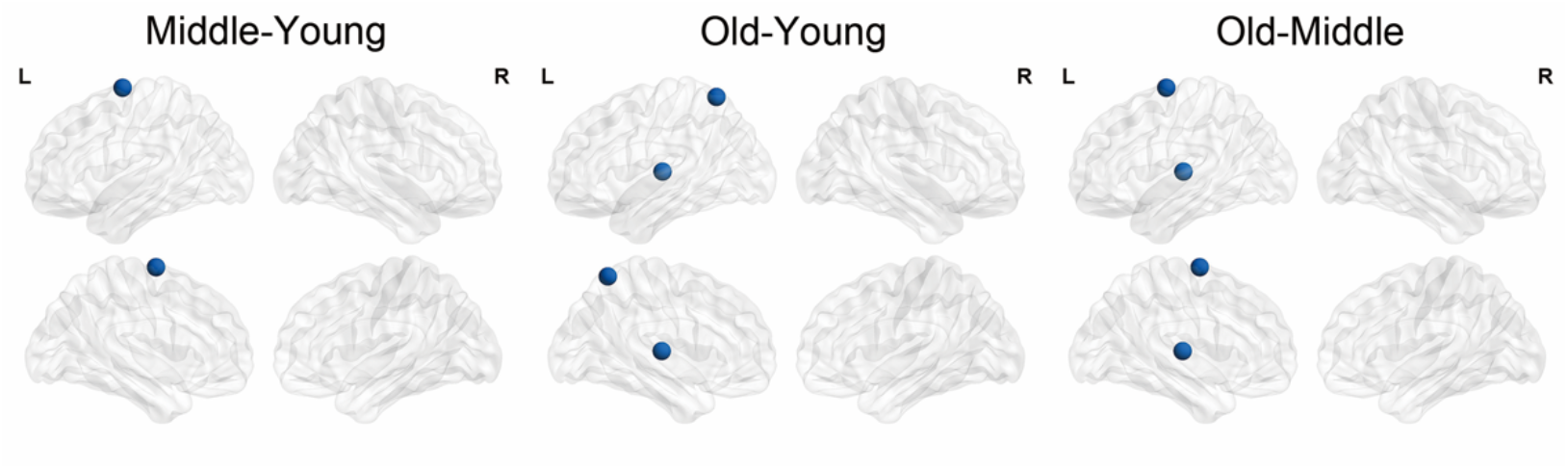
Between-group differences of the nodal overlapping ratio. The dark blue node indicates significant between-group difference *(P* < 0.001, 10000 permutation).

**Figure 6.**
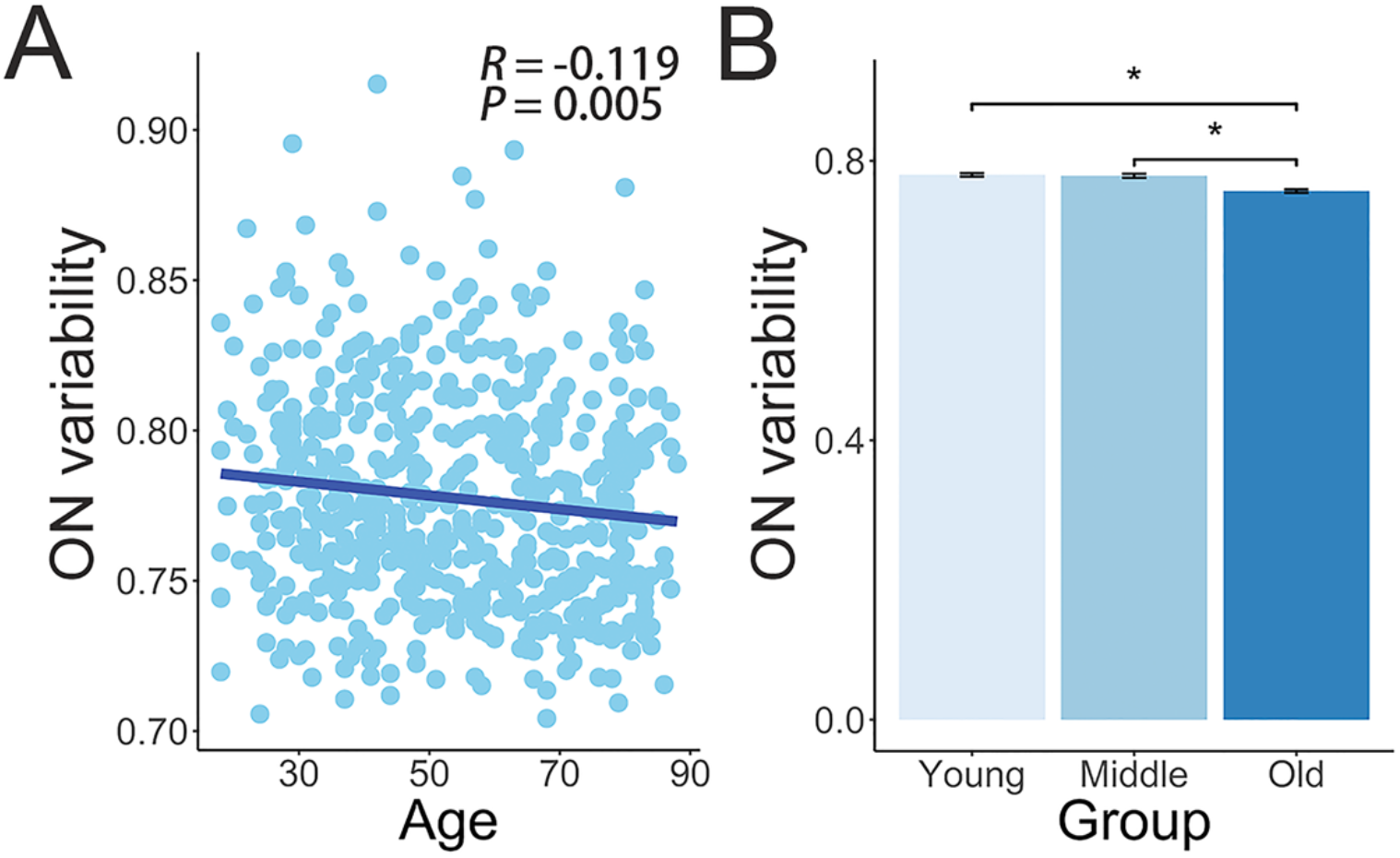
Lifespan changes in overlapping nodes (ON) regarding (A-B) variability of spatial distribution. The light blue nodes denote participants, and dark blue lines denote the aging regression line for linear model. In bar plot, the asterisk indicates significant between-group difference *(P* < 0.05, 10000 permutations, Bonferroni-corrected).

### 3.3 Adult lifespan changes in the functional characteristics of overlapping nodes

The nodal overlapping probability was found positively correlated with the gradient (*R* = 0.397, *P* < 10^−3^; Figure 7A), and this positive correlation maintained within each age group (Young, *R* = 0.362, *P* < 10^−3^; Middle, *R* = 0.416, *P* < 10^−3^; Old: *R* = 0.339, *P* < 10^−3^; Figure 7B). In addition, the nodal overlapping probability was also found positively correlated with the functional flexibility (*R* = 0.401, *P* < 10^−3^; Figure 7C). The positive correlations also maintained within each age group (Young, *R* = 0.397, *P* < 10^−3^; Middle, *R* = 0.324, *P* < 10^−3^; Old, *R* = 0.394, *P* < 10^−3^; Figure 7D). These results revealed that the overlapping nodes tended to have higher functional gradient and flexibility.

**Figure 7.**
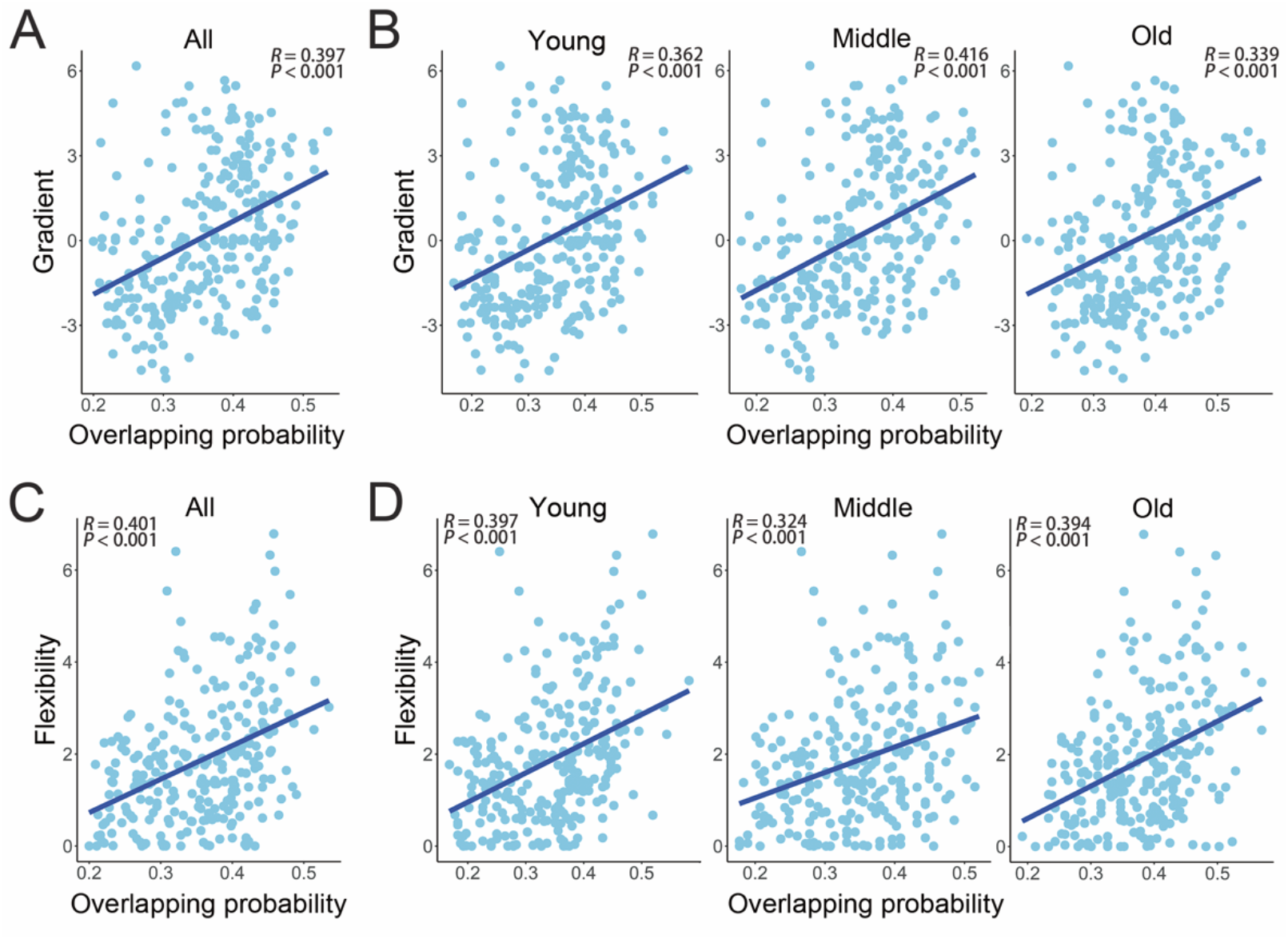
Lifespan changes in the function al characteristics of overlapping nodes (ON) overlapping probability functional characteristics regarding (A & B) gradient 1, (C & D) functional flexibility. The light blue nodes denote participants, and dark blue lines denote the aging regression line for linear model.

### 3.4 Relationships between characteristics of overlapping modules or nodes and cognitive performances

For fluid intelligence, we found that it had positive correlations with the overlapping modularity (*R* = 0.167, *P* < 10^−3^), the modular similarity (*R* = 0.367, *P* < 10^−3^), the individual variability in the spatial pattern of overlapping nodes (*R* = 0.088, *P* = 0.039), and the modular overlapping percentage in CON and SAN (CON: *R* = 0.089, *P* = 0.037; SAN: *R* = 0.098, *P* = 0.021, Figure 8A). In contrast, we found that fluid intelligence had negative correlations with the number of overlapping nodes (*R* = −0.134, *P* = 0.002), the number of overlapping nodes that participated in both two and three overlapping modules (*k* = 2: *R* = −0.110, *P* = 0.010; *k* = 3: *R* = −0.138, *P* = 0.001), and the modular overlapping percentage in VIS (*R* = −0.156, *P* < 10^−3^, Figure 8B). As for the Benton face recognition test score, similar as the patterns of fluid intelligence, it was found positively correlated with the overlapping modularity (*R* = 0.149, *P* < 10^−3^) and the modular similarity (*R* = 0.252, *P* < 10^−3^, Figure 8C-D), but negatively correlated with the number of overlapping nodes (*R* = −0.091, *P* = 0.034), the number of overlapping nodes that participated in both three or more overlapping modules (*k* = 3: *R* = −0.106, *P* = 0.010; *k* ≥ 4: *R* = −0.098, *P* = 0.022) and the modular overlapping percentage in VIS (*R* = −0.148, *P* < 10^−3^, Figure 8E).

**Figure 8.**
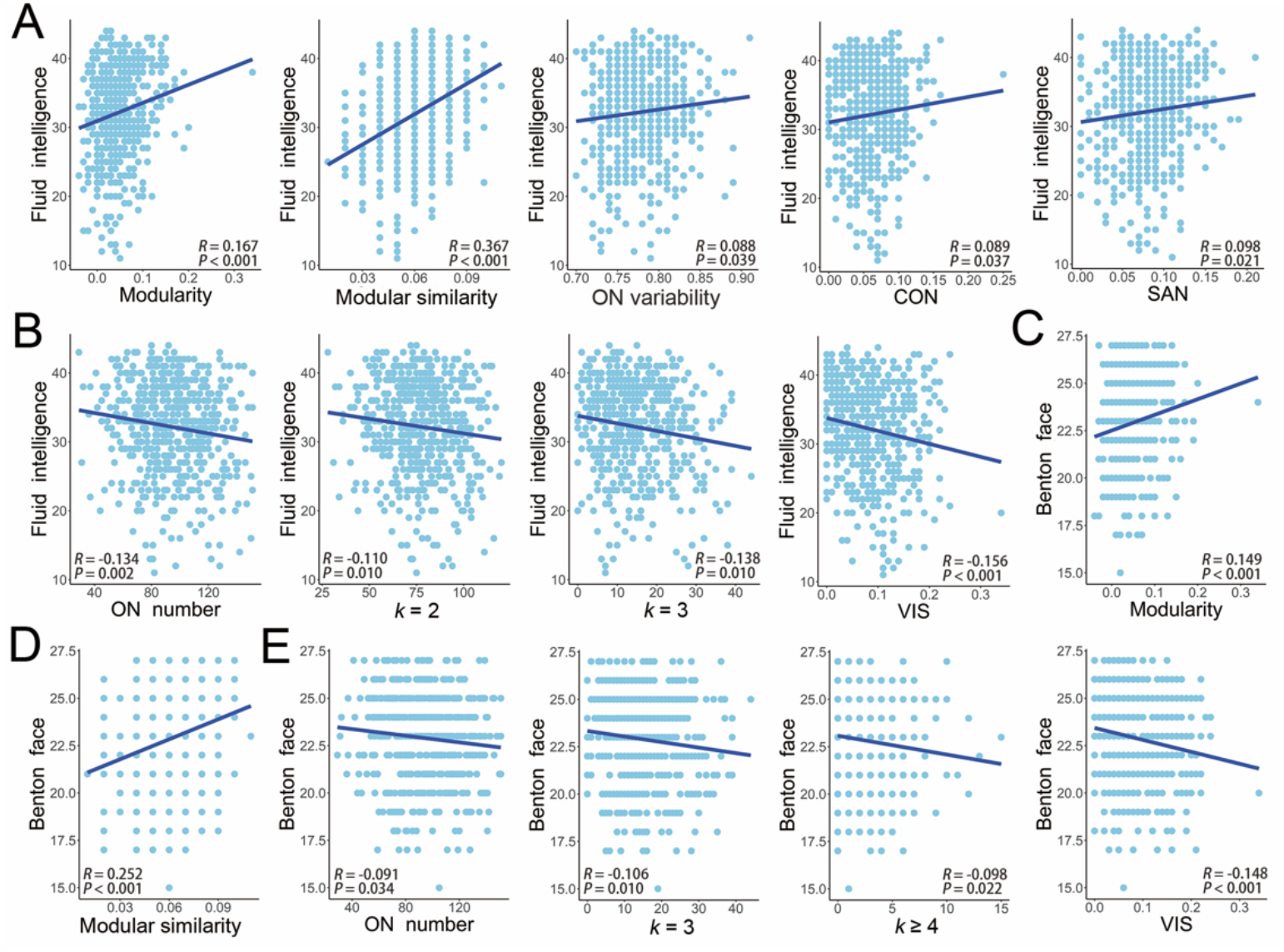
Relationship between characteristics of overlapping modules (OM) or nodes (ON) and cognitive performances including (A & B) fluid intelligence and (C, D & E) Benton face recognition. The light blue nodes denote participants, and dark blue lines denote the aging regression line for linear model.

Additionally, we found that the decrease in the overlapping modular similarity was associated with age (path a: β = −0.436, *P* < 10^−3^). After controlling the influence of age, the higher overlapping modular similarity was related to higher fluid intelligence (path b: β = 0.092, *P* < 10^−3^). After considering the effect of overlapping modular similarity, the effect of age on fluid intelligence was weakened (path c’: β = −0.629, *P* < 10^−3^, from path c: β = −0.669, *P* < 10^−3^, Figure 9A). This mediation analysis revealed that overlapping modular similarity was a significant mediator (indirect effect = −0.040, 95% CI = [−0.068, −0.013]) and could partially explain the negative association between age and fluid intelligence (Figure 9A). Besides, we found that the increase of the modular overlapping percentage in VIS was associated with age (path a: β = 0.149, *P* < 10^−3^). After controlling the influence of age, the higher modular overlapping percentage in VIS was related to the lower Benton face recognition test score (path b: β = −0.077, *P* < 10^−3^). After considering the effect of modular overlapping percentage in VIS, the effect of age on Benton face recognition test score was weakened (path c’: β = −0.473, *P* < 10^−3^, from path c: β = −0.484, *P* < 10^−3^, Figure 9B). This mediation analysis revealed that modular overlapping percentage in VIS was a significant mediator (indirect effect = −0.011, 95% CI = [−0.026, −0.001]) and could partially explain the negative association between age and Benton face recognition (Figure 9B).

**Figure 9.**
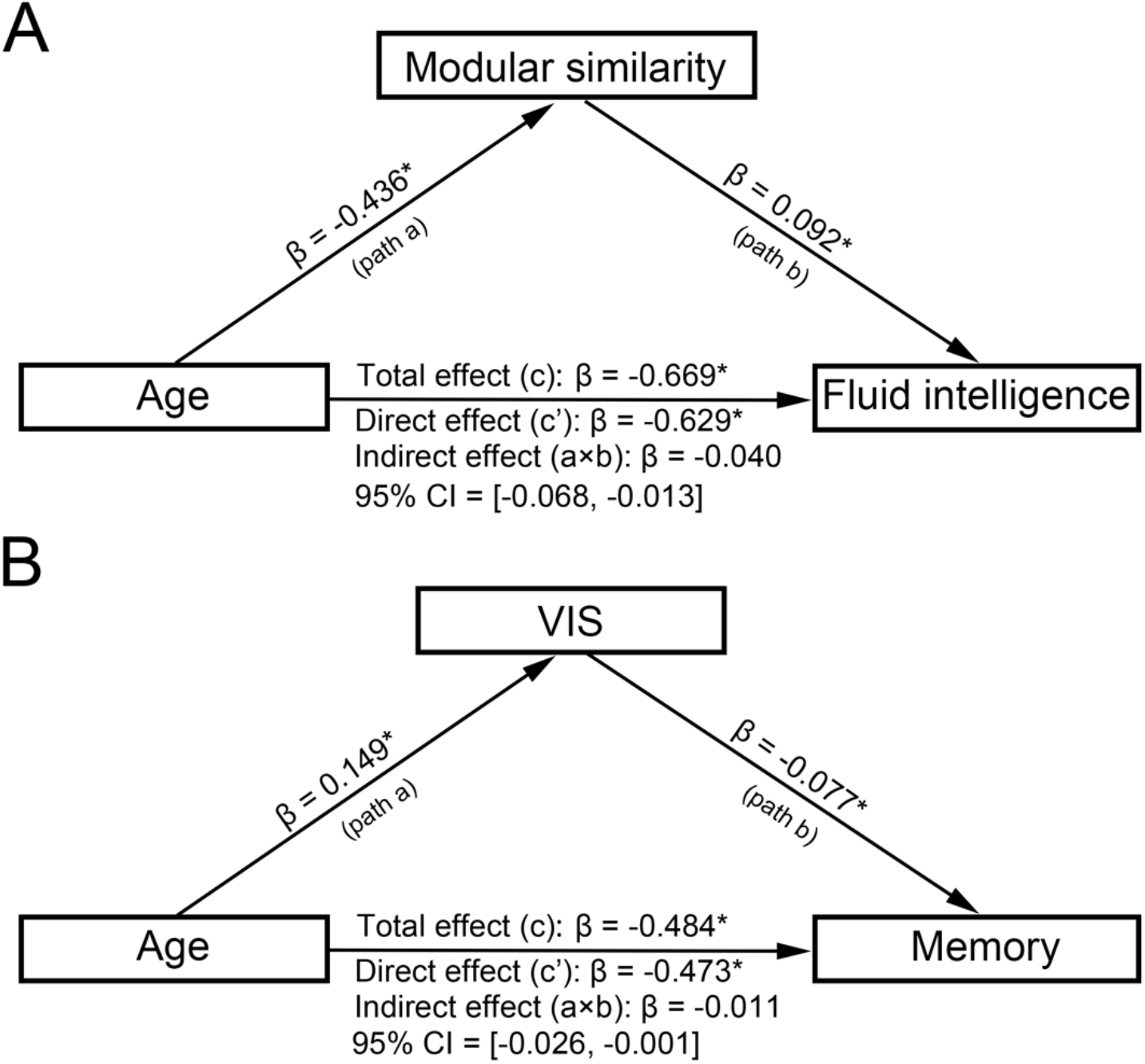
Mediating effects of overlapping module or node characteristics on lifespan changes in (A) fluid intelligence and (B) memory stands for Benton face recognition test performance. Standardized regression coefficients were reported, and the asterisk indicates significant relationship.

## 4. Discussion

In this study, we studied the change of the overlapping functional modular organization across the adult lifespan. In terms of overlapping functional modules, the overlapping modularity and the modular similarity both gradually declined linearly over the adult lifespan. In terms of the overlapping nodes, the number increased linearly with age. As for their distribution, the modular overlapping percentage linearly increased in VIS but decreased in FPN with age. Besides, the individual variability in the spatial distribution of overlapping nodes linearly decreased over the adult lifespan. Both the nodal functional gradient and flexibility were positively correlated with the nodal overlapping probability during the adult lifespan and in each age group. Finally, the age-related characteristics of overlapping modules and overlapping nodes were found correlated with memory-related cognitive performance. Together, our results indicated lifespan decreased segregation of the brain functional modular organization, providing new insight into the age-related changes in brain function and behavioral performance.

### 4.1 Adult lifespan changes of overlapping modules

Modularity, which measures how well a network can be decomposed into a set of sparsely inter-connected but densely intra-connected modules (Newman, 2004), is an advanced topological property of brain network organization and can be used to evaluate the functional segregation (Meunier, Lambiotte, Fornito, Ersche, & Bullmore, 2009). The present study found that the overlapping modularity of the brain functional network linearly decreased over the adult lifespan, which is consistent with the findings in non-overlapping modularity studies (Betzel et al., 2014; Cao et al., 2014; Geerligs et al., 2014). These consistent findings of declined modularity in the elderly revealed that aged brains had decreased intra-module functional connectivity and increased inter-module functional connectivity, which suggests the brain functional network of elder people to be less segregated or less differentiated (Betzel et al., 2014; Chan et al., 2014; Geerligs et al., 2014). The dedifferentiation theory suggests that overactivation of brain regions in the elderly during cognitive tasks may be caused by the decrease in functional distinction between regions (Baltes & Lindenberger, 1997; Park et al., 2004), which may further induce the functional module-level dedifferentiation, as observed in the current study. Our findings provided direct support for the dedifferentiation phenomenon at the functional module level, which could also account for the less distinctive neural representations in old age (Li, Lindenberger, & Sikström, 2001).

### 4.2 Adult lifespan changes of overlapping nodes

During the adult lifespan, regions in high-order cognitive functions were found to have high overlapping probabilities. Additionally, we found that the regions with low nodal overlapping probabilities were mostly involved in primary functional modules. Thus, these results suggest that higher-order associative modules are more likely to embed overlapping nodes, which was largely compatible with previous overlapping module studies in healthy young adults (Lin et al., 2018; Yeo et al., 2014). Notably, a similar nodal overlapping probabilities distribution existed over the adult lifespan, suggesting relative preservation of the crucial roles of these regions.

Besides the distribution of overlapping nodes, we also examined how the characteristics of overlapping nodes changed during the adult lifespan. We found a significant linear increase of the overlapping node number during the adult lifespan. Additionally, the overlapping probability of the region in the left thalamus was higher in the older group than in the other groups, indicating that this region was more likely to participate in multiple functional modules in elder people. The thalamus is involved in multiple cognitive functions, which is an integrative hub for functional brain networks (Hwang, Bertolero, Liu, & D’esposito, 2017). Previous studies have found that older adults exhibited stronger functional connectivity between the thalamus and putamen, which highlights the potential role of enhanced thalamic connectivity in protecting the memory ability from aging (Ystad, Eichele, Lundervold, & Lundervold, 2010). Also, the region in the left superior parietal lobule had higher overlapping probabilities in the old group than in the young group, which was also associated with working memory (Jager, Kahn, Van Den Brink, Van Ree, & Ramsey, 2006). Thus, we speculated that regions with higher overlapping probabilities in the elderly might be the compensation for the age-related decrease in functional connectivity.The results on the functional characteristics showed that the regions with higher nodal overlapping probability tended to have higher gradient value and higher flexibility during the adult lifespan. This positive relationship was also confirmed in each age group. It also implies the importance of analyzing overlapping structures for understanding brain functional changes during the adult lifespan and across age groups.

In general, we found that the indicators capturing decreased characteristics of overlapping modules and overlapping nodes during the adult lifespan (e.g., modularity, individual variability in the overlapping node distribution) tended to have positive correlations with fluid intelligence and the Benton face recognition scores. Conversely, the indicators capturing increased characteristics of overlapping modules and overlapping nodes during the adult lifespan (e.g., number and diversity of overlapping nodes) tended to have negative correlations. Combining these results with previous findings that fluid intelligence and the Benton face recognition test score was positively related to memory performance (Benton et al., 1994; Cattell, 1971) and declined with age (Feng et al., 2020; Kievit et al., 2014), we speculated that the age-related changes in overlapping modules and overlapping nodes were closely associated with the age-accompanied decline in memory ability. In addition, we found that the overlapping modular similarity and the overlapping percentage of VIS partially mediated the negative associations of age with fluid intelligence and the Benton face recognition test score, respectively. Besides, the mediation effect of overlapping modular similarity on lifespan changes in fluid intelligence was maintained in most validation results. Thus, the overlapping modular similarity might serve as a biomarker for aging. Together, these results further supported our conjecture that the age-related changes in the overlapping functional modular organization were possible neural representations of cognitive performance change across the lifespan.

### 4.3 Future consideration

Several methodological limitations need further considerations. First, the functional atlas used in our study was obtained by exploring a combination of meta-analysis of functional connectivity in an adult population (Power et al., 2011). Ideally, individual functional atlas should be determined by each participant-specific functional connectivity for individual analysis. Several studies have focused on the methodology of individualized functional atlas (Cui et al., 2020; Kong et al., 2019; Wang et al., 2015). However, there is still not yet a uniformly recognized golden method about the individualized functional atlas. Future improvement in atlas-related methodological research may help us to further validate and deepen our findings based on reliable individualized functional atlas. Second, whether the global signals should be removed is currently still debatable in the preprocessing procedure of the R-fMRI. Both our study and previous ones have found large differences between results with and without regressing out the global signals (Li et al., 2016; Murphy et al., 2009; Yang et al., 2014). Future studies are necessary to propose better approaches to minimize the noise in R-fMRI and to evaluate the optimal approaches to analyze the global signals. Last, although the current research dataset already covers a wide age range, it is still not a longitudinal/follow-up dataset that covers the whole adult lifespan of each participant. In the future, further validation based on true follow-up data should be considered.

## Notes

### Competing Interest Statement

The authors have declared no competing interest.

